# Quantitative Models Identify Histone Signatures of Poised Genes Prior to Cellular Differentiation

**DOI:** 10.1101/137646

**Authors:** Rui Tian

## Abstract

**Background:** Recent studies have shown that histone marks are involved in pre-programming gene fates during cellular differentiation. Bivalent domains (marked by both H3K4me3 and H3K27me3) have been proposed to act in the histone pre-patterning of poised genes; however, bivalent genes could also resolve into monovalent silenced states during differentiation. Thus, the histone signatures of poised genes need to be more precisely characterized.

**Results:** Using a support vector machine (SVM), we show that the collective histone modification data from human blood hematopoietic cells (HSCs) could predict poised genes with fairly high predictive accuracy within the model of directed erythrocyte differentiation from HSCs. Surprisingly, models with single histone marks (e.g., H3K4me3 or H2A.Z) could reach comparable predictive powers to the full model built with all of the nine histone marks. We also derived an H2A.Z and H3K9me3-based Naive Bayesian model for inferring poised genes, and the validity of this model was supported by data from several other pluripotent/multipotent cells (including mouse ES cells).

**Conclusion:** Our work represents a systematic quantitative study that verified that histone marks play a role in pre-programming the activation or repression of specific genes during cellular differentiation. Our results suggest that the relative quantities of H2A.Z modification and H3K9me3 modification are important in determining a corresponding gene’s fate during cellular differentiation.

## Background

The tremendous advances in genomic sequencing have allowed access to an unprecedented plethora of genetic data, i.e., DNA sequences. Yet, we are still far from understanding global transcriptional regulation in eukaryotic cells under normal conditions, let alone in disease states. Gene transcriptional regulation at the epigenetic level modulates the complex process by which a single genome (e.g., a zygote) can develop into one of over 200 types of different cells, which collectively form the various tissues and organs of a human body [1]. Indeed, the versatility of epigenetic regulation at the organismal level is remarkable.

Epigenetic regulation is mainly achieved *via* DNA methylation and histone modifications [2]. Histone N terminal tails, when chemically modified, e.g., methylated, acetylated, or phosphorylated (also referred to as histone marks), are closely associated with gene transcription in either a positive or negative manner. The intensively studied histone marks include H3K4me3, H3K27me3, H3K36me3 and H3K4me1 [3-7]. Bernstein et al. (2006) described bivalent domains in mouse embryonic stem cells (mES cells), referring to those genomic regions with both H3K4me3 (an activating histone mark) and H3K27me3 (a repressive histone mark) [8]. These authors proposed that the genes harboring bivalent domains (hereafter referred to as bivalent genes) in stem cells are repressed but poised for expression as the cells differentiate. Recently, there have also been reports showing that histone marks other than H3K4me3 and H3K27me3 may also play a pre-programming role in development and that histone mark pre-patterns may function similarly even in lineage-restricted multipotent cells [9, 10].

To date, previous studies characterizing the pre-programming role of histone marks have been largely empirical. Several reports (including Bernstein et al. in 2006) have demonstrated that the fate of bivalent genes is not necessarily activation: certain genes can be repressed [11, 12]. Therefore, it is difficult to predict whether a gene is poised for expression by simply observing whether it has both H3K4me3 and H3K27me3 modifications. Accordingly, the histone modification feature of poised genes within the context of cellular differentiation requires a more precise characterization. In addition, bivalent genes are also present in tissue stem cells or other terminally differentiated cells and can be generated *de novo* [11, 13, 14]. Thus, it remains to be explored whether a common histone signature is present for all poised genes, regardless of cell identity or species.

Using a support vector machine (SVM), we demonstrate that within the model of directed erythrocyte differentiation from human blood hematopoietic cells (HSCs) [11], the collective histone modification data from HSCs could predict induced (poised) genes with fairly high predictive accuracy. Surprisingly, even models with a single histone mark (e.g., H3K4me3 or H2A.Z) exhibited a predictive power that was comparable to the full model using all nine histone marks. We then derived an H2A.Z and H3K9me3-based Naive Bayesian model for inferring poised genes, and the validity of this model was supported by data from several other pluripotent/multipotent cells (including mouse ES cells). Our results further verified the pre-programming role of histone marks and suggested that relative quantities of H2A.Z and H3K9me3 modifications are important in determining a corresponding gene’s fate during cellular differentiation.

## Results

### The predictive role of HSC histone marks for the activation of lineage-specific genes in erythrocyte precursors

Cui et al. (2009) [11] generated a set of histone modification (9 types) ChIP-seq and microarray data from HSCs and erythrocyte precursors differentiated from HSCs. As both the progenitor and descendant cells share the same genetic composition and the transcriptomic and epigenomic data for both are available, the Cui et al. (2009) data represent a suitable resource for investigating the histone modification features of poised genes within the context of (tissue-) stem cell differentiation and are hereafter referred to as the HSC dataset. First, the raw microarray data for both cell types were analyzed using the MAS5.0 P/M/A calling algorithm (see Materials and Methods for details). In total, 369 genes (RefSeq transcripts) were identified as silent in HSCs but active in erythrocyte precursors; these data are hereafter referred to as the poised gene set. A total of 8,007 genes (RefSeq transcripts) were consistently silent in both cell types; these data are hereafter referred to as the silenced gene set. The histone modification signal densities covering the promoter and gene body regions were calculated separately for these two sets of genes (detailed in the Materials and Methods). Pairwise comparisons between the two sets of genes demonstrated that they largely differed for many histone marks in terms of their signal densities (in additional file, Figure S1). In general, poised genes are characterized by higher signal levels of active marks and lower signal levels of repressive marks, suggesting that poised and silenced genes are distinctly marked in progenitor cells (in this case, HSCs) [11].

Overall, poised genes differ substantially from silenced genes in their histone profiles; however, it is difficult to determine individually whether a gene is poised based only on a single histone mark. For example, as is shown in Supplementary Figure S1, a large proportion of poised genes overlap with silenced genes in terms of their H2A.Z signal levels. Yet, when the H3K4me3 signal levels of the two sets of genes are plotted against the corresponding H3K27me3 signal levels, it is difficult to draw a simple curve to discriminate the two sets of genes (in additional file, Figure S2). This is also true when the information of H3K36me3 is incorporated by visualizing its signal levels using different dot sizes in the same scatter plot. Furthermore, as shown in Figure S2, some genes, either belonging to the poised or silenced gene set, are characterized by a very low level of H3K4me3 but a very high level of H3K36me3 (accounting for 9.5% of the poised gene set and 3.2% of the silenced gene set). Because H3K36me3 is a well-established histone mark associated with transcriptional elongation and transcription of a gene normally occurs only when H3K4me3 is present at its promoter, we did not include these genes for further analysis.

To ascertain whether a given gene is poised based on its histone mark information, we set out to build a support vector machine (SVM) classifier. In this study, we initially considered the histone modification ChIP-seq read densities for both the promoter and transcribed regions, as it was unclear which region is more informative for a particular histone mark. We then calculated the F score for each feature using the formula presented in the Materials and Methods. F score is viewed as a classifier-independent measure of discriminant ability of a feature to separate two classes[15]. It is demonstrated in Figure 1a that for H2A.Z, H3K27me3, H3K4me3, H3K9me3, the features based on promoter regions are more informative; One the contrary, for the 4 mono-methylation histone marks (i.e., H3K27me1, H3K4me1, H3K9me1 and H4K20me1), the features based on gene body regions are more informative.

**Figure 1.**
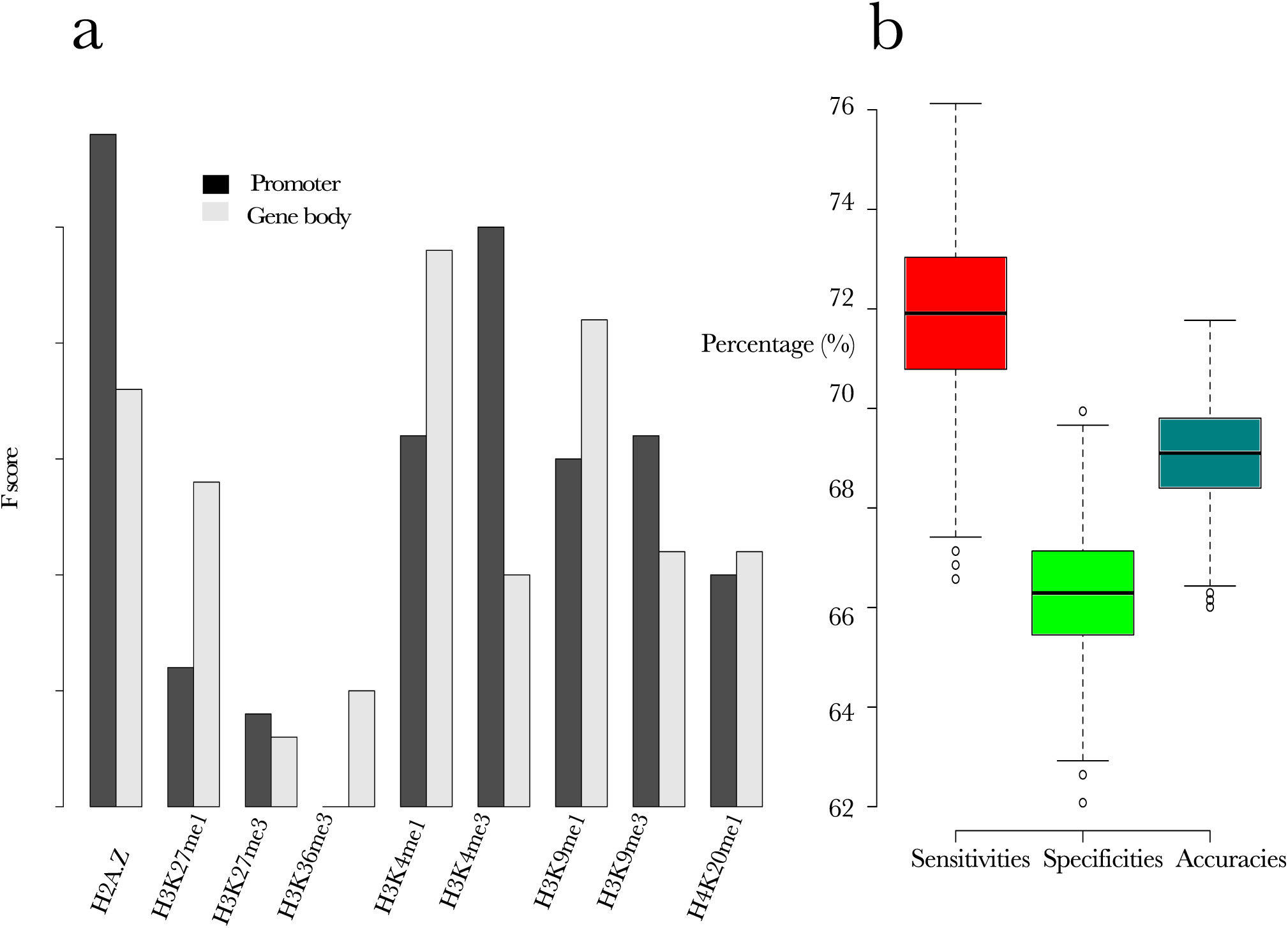
Histone marks in HSCs are predictive of poised genes. The panel (a) illustrates the discriminant ability of each histone mark feature as measured by calculation of F score (detailed in Materials and Methods section). The panel (b) shows the statistics of 2, 000 SVM models’ performances. Sensitivity refers to true positive rate; specificity refers to true negative rate; accuracy is the rate of correct predictions among the total predictions.

Based on these observations, for each histone mark only the more informative feature was retained for the subsequent model building process. The training and test of the SVM classifiers was implemented in the R computing environment using the package “e1071” [16](details are in the Materials and Methods). Because there are approximately 20-fold more negative instances (i.e., silenced genes) than positive instances (i.e., poised genes), the positive instances were under-sampled to an amount of instances that was equal to that of the positive instances. The under-sampled negative instances were then combined with the original positive instances for building the models. The predictive power was determined using a five-fold cross-validation method, and the process was repeated more than 2,000 times. As shown in Figure 1b, the median of the sensitivities, specificities and accuracies are 71.9%, 66.3%, and 69.1%. These good performances indicate that the model is, indeed, applicable to infer whether a specific gene is poised for subsequent activation during cellular differentiation from the histone modification status of the progenitor cell.

### Different discriminative roles of single histone marks for poised genes

We have shown that one can predict whether a gene is poised using nine histone mark features in a framework of SVM learning. It has been shown previously that many histone marks are closely correlated to each other and that, in a sense, different histone marks may carry redundant information with regard to gene regulation and chromatin states [7]. As shown in Figure S3 (in additional file), our correlation analysis is highly consistent with the previous findings and identified several closely correlated histone mark pairs or groups [4]. For example, the active marks H2AZ, H3K27me1, H3K4me1, H3K4me3, H3K9me1 and H3K36me3 are highly correlated to each other, and these active marks generally show a weak negative correlation with the repressive marks H3K27me3 and H3K9me3. Interestingly, H3K27me3 is dramatically negatively correlated with H3K27me1; conversely, the negative correlation between H3K9me3 and H3K9me1 is not very pronounced.

To define clearly the relative contribution of a certain histone mark feature to the prediction model, it is necessary to rule out the redundant information of highly correlated histone marks. The SVM prediction models were rebuilt using a single histone mark feature each time. The overall performances for more than 2,000 sampled subsets of data (the procedure was the same as described above) were compared with the original trained model based on all of the nine-histone mark features (referred to as the full model) (Figure 2). Surprisingly, when each was used alone to train a prediction model, three of the histone features (H3K4me1, H3K9me1 and H4K20me1), showed even slightly higher sensitivities than those of the full model but at the sacrifice of much compromised specificities. In contrast, prediction models based on H3K27me1 and H3K36me3 separately exhibited high specificities at the cost of much lower sensitivities. Only models trained based on H3K4me3 and H2A.Z separately reached balanced sensitivities and specificities that were comparable (but slightly lower) to those of the full model. Interestingly, H3K27me3, when used to train prediction models alone, was barely effective.

**Figure 2.**
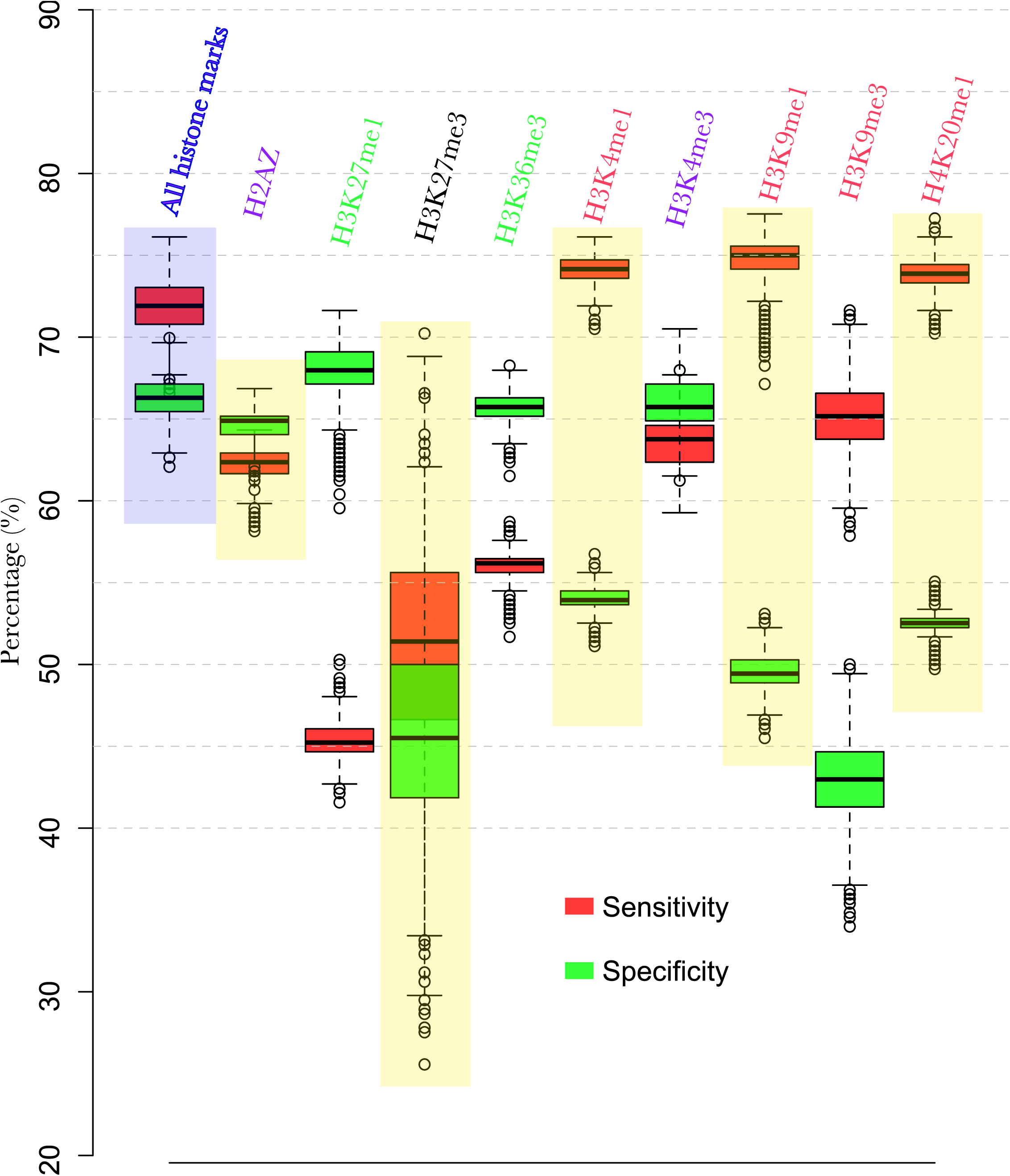
Single histone marks vary in discriminant capabilities for poised genes.

Based on these results, the histone marks can be classified into three major categories: Class i, those that can reach high sensitivities with compromised specificities (H3K4me1, H3K9me1 and H4K20me1); Class ii, those that can reach high specificities with compromised sensitivities (H3K27me1 and H3K36me3); and Class iii, those that can reach both high sensitivities and specificities (H3K4me3 and H2A.Z) and can be regarded concurrently as Class i or Class ii histone marks. The performance differences of these investigated single-feature prediction models suggest that some of the histone marks are more efficient in identifying a certain gene as a poised gene (Class i histone marks, e.g., H3K4me1), whereas others are more efficient in identifying that a certain gene is not a poised gene (Class ii histone marks, e.g., H3K36me3). H3K4me3 and H2AZ are exceptional, as they are both good predictors for identifying potential poised genes and ruling out silenced genes. We also compared the predictive performances of triple histone marks by enumerating all of the possibilities and found that a preferable triple model should comprise at least one Class i histone mark and at least one Class ii histone mark (data not shown).

### A two-histone mark probabilistic model for predicting poised genes in various cell types

Using the SVM, we showed that the quantitative histone mark models in progenitor cells are predictive of poised genes, and we also assessed the predictive powers of each single histone mark. Nevertheless, although powerful, SVM models have been criticized for their “black-box” nature. Thus, to derive an intuitive and simplified probabilistic model, we investigated the dependency relationships among “gene poising” and histone mark features using the WinMine Tookit [17, 18]. The histone profiles for the total RefSeq genes were discretized by k-means clustering into three levels (“Lev0”, “Lev1”, and “Lev2”, from the lowest to highest levels). Because the training dataset of the HSCs [11] is severely unbalanced, under-sampling was performed in each case to obtain a random subset of poised genes (250 genes) and a random subset of silenced genes (250 genes), and the combined instances were used for the learning Bayesian Networks. To obtain robust and reproducible Bayesian Networks, the aforementioned process was repeated 100 times, and only the edges directed to “gene poising” that appeared more than 30 times were retained. Among the 100 learned Bayesian networks, 47% contained two edges pointing to gene poising, 26% contained three edges pointing to gene poising, and the left contained more edges. For simplicity, we chose a two-histone mark model (namely H2AZ+H3K9me3), as shown in Figure 3a.

**Figure 3.**
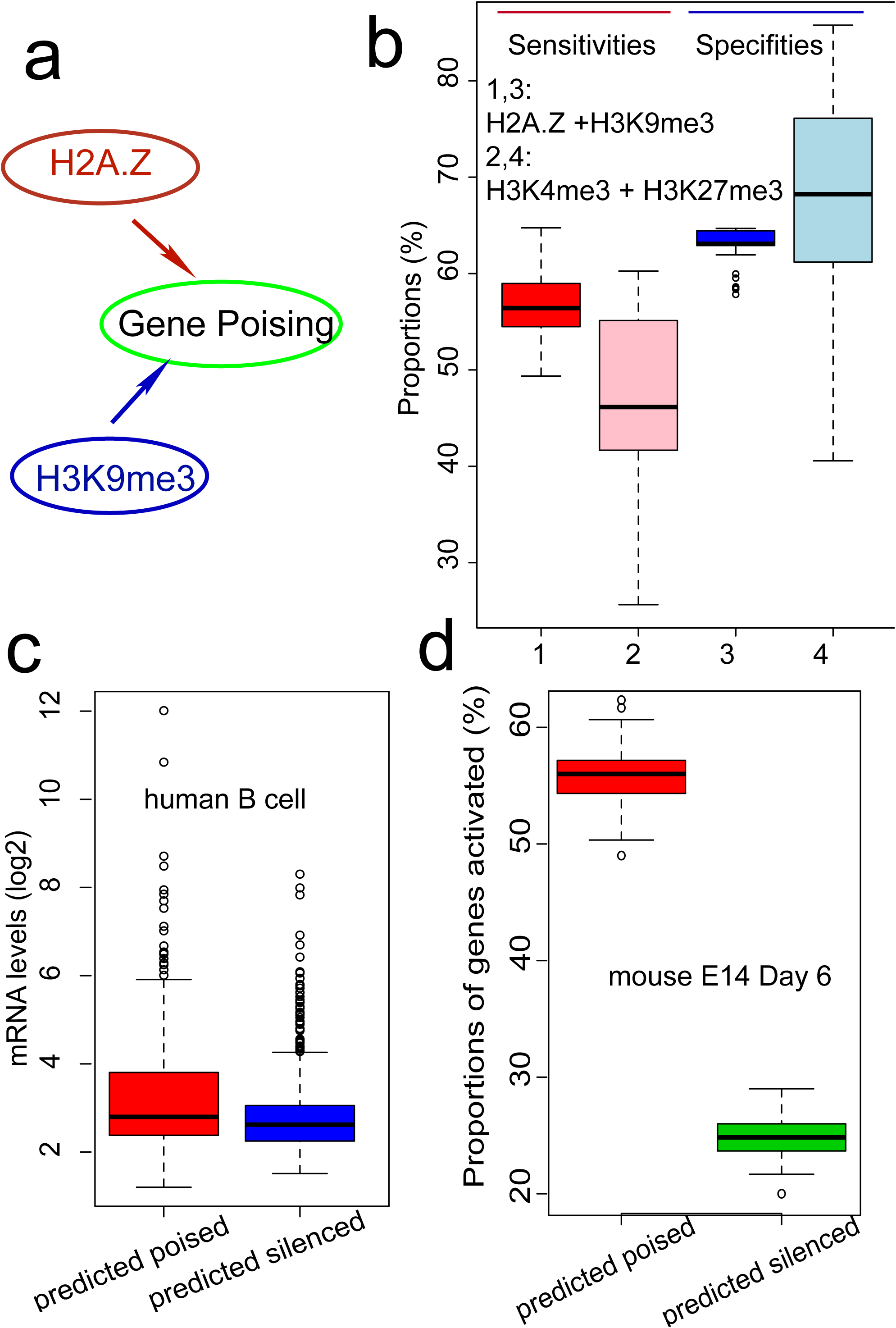
H2A.Z and H3K9me3 Naïve Bayesian models distinguish poised genes from silenced genes in multiple cell lines. The panel (a) compares the performances of H2A.Z+ H3K9me3 models between H3K4me4 + H3K27me3 models by repeating 100 times. The panel (b) shows the selected Bayesian network learned from the HSC dataset, where only directed edges pointing to gene poising are illustrated. A red arrow indicates a positive role while blue one indicates a negative role. The panel (c) compares the log2 value of expression levels of predicted poised genes versus silenced genes in B cells. The panel (d) demonstrates that predicted poised genes in mouse ES cells are almost twice likely to be activated upon 6 days induction compared with predicted silenced genes.

According to Bayesian network theory, a node is only dependent on the directed edges pointing to it. Thus, if our learned Bayesian network is reliable, we should be able to deduce “gene poising” solely from the profile data of the two-histone marks (H2AZ+H3K9me3). For simplification of the probabilistic model, we assumed independence between H2AZ and H3K9me3 profiles and derived Naive Bayesian models using the HSC training data. The models were repeatedly learned using randomly sampled 200 poised plus 200 silenced genes and validated using the rest of the data. As shown in Figure 3b, the median sensitivity is approximately 56.4% and the median specificity is approximately 63.1%. As Bernstein et al. (2006) [8] proposed the co-existence of H3K4me3 and H3K27me3 as an indictor for poised genes, we also tested the predictive power of an H3K4me3+H3K27me3 Naive Bayesian model. In general, our H2AZ+H3K9me3 model is more stable than the H3K4me3+H3K27me3 model, though the former shows a lower specificity.

To test whether the H2AZ+H3K9me3 Naïve Bayesian model learned from the HSC dataset is also valid in other cells of variable differentiation potentials, we selected an optimal model from 100 learned models and applied it to GM12878 cells, a lymphoblastoid cell line that may represent B cell progenitor cells. Transcriptionally inactive genes were first selected by microarray analysis, and the log likelihood ratio s was calculated for each of these silent genes (detailed in the Materials and Methods: the higher is the *s* value, the more likely it is that the corresponding gene is poised). Because the prediction model showed a modest sensitivity and specificity, to test whether the prediction was reasonable, only the genes with the highest s scores were predicted to be poised genes and the genes with the lowest s scores were predicted to be silenced. In the GM12878 cells, 240 ENSEMBL genes were predicted to be poised, whereas 527 ENSEMBL genes were predicted to be silenced. By comparing the expression values of these two groups of genes in B cell (biogps database, http://biogps.org/), it was found that there were significantly more genes of the predicted poised gene group that were activated in B cells (Fig 3c). This result suggests that our H2AZ+H3K9me3 model learned from the HSC dataset could also be applied to other cells (e.g., B lymphoblastoid cells).

We also tested the H2AZ+H3K9me3 Naïve Bayesian model with mouse embryonic stem E14 cells. The GSE36114 dataset contains E14 ChIP-seq and RNA-seq data of E14 cells at day 0 and day 6 (upon differentiation). Using the model trained with the HSC dataset, poised (2,577) and silenced (556) genes were predicted with a high confidence based on the E14 cell day 0 histone modification data. As there were many more genes predicted to be poised than silenced genes, repeated under-sampling (300 genes from each group) was performed to compare the ratio of genuine activated genes (day 6) in these two groups. As illustrated in Figure 3d, the predicted poised genes were, on average, almost twice as likely to be activated during differentiation. This result indicates that our H2AZ+H3K9me3 Naïve Bayesian model learned from an HSC dataset (tissue stem cells) can also successfully be applied to mouse ES cells, thereby suggesting a common poising mechanism by histone marks, at least in mammalian cells.

### Histone modification features of poised genes

Based on our H2A.Z+H3K9me3 Naïve Bayes model trained with the HSC dataset, we proposed to calculate the log likelihood ratio of each gene to be poised versus silenced. As each histone feature was discretized into three levels, there would be a total of nine types of combinations of varying H2A.Z and H3K9me3 levels. When the score s is ordered according to the ascending levels of each histone mark, an apparent trend about the likelihood of the gene to be poised is observed (Figure 4a). Genes with higher levels of H2A.Z plus lower levels of H3K9me3 are more likely to be poised, whereas genes with lower levels of H2A.Z plus higher levels of H3K9me3 are more likely to be silenced. Figure 4b shows the histone modification profiles of two genes (one is poised, the other is silenced) in HSCs. Although these two genes appear to both be bivalent (marked by both H3K4me3 and H3K27me3), a closer look indicates that the silenced gene displays stronger in both H3K9me3 and H3K27me3 signals and that its H2A.Z signal is weaker than that of the poised gene. These two examples also show the validity of our method for capturing the information of histone marks by focusing on promoter regions.

**Figure 4.**
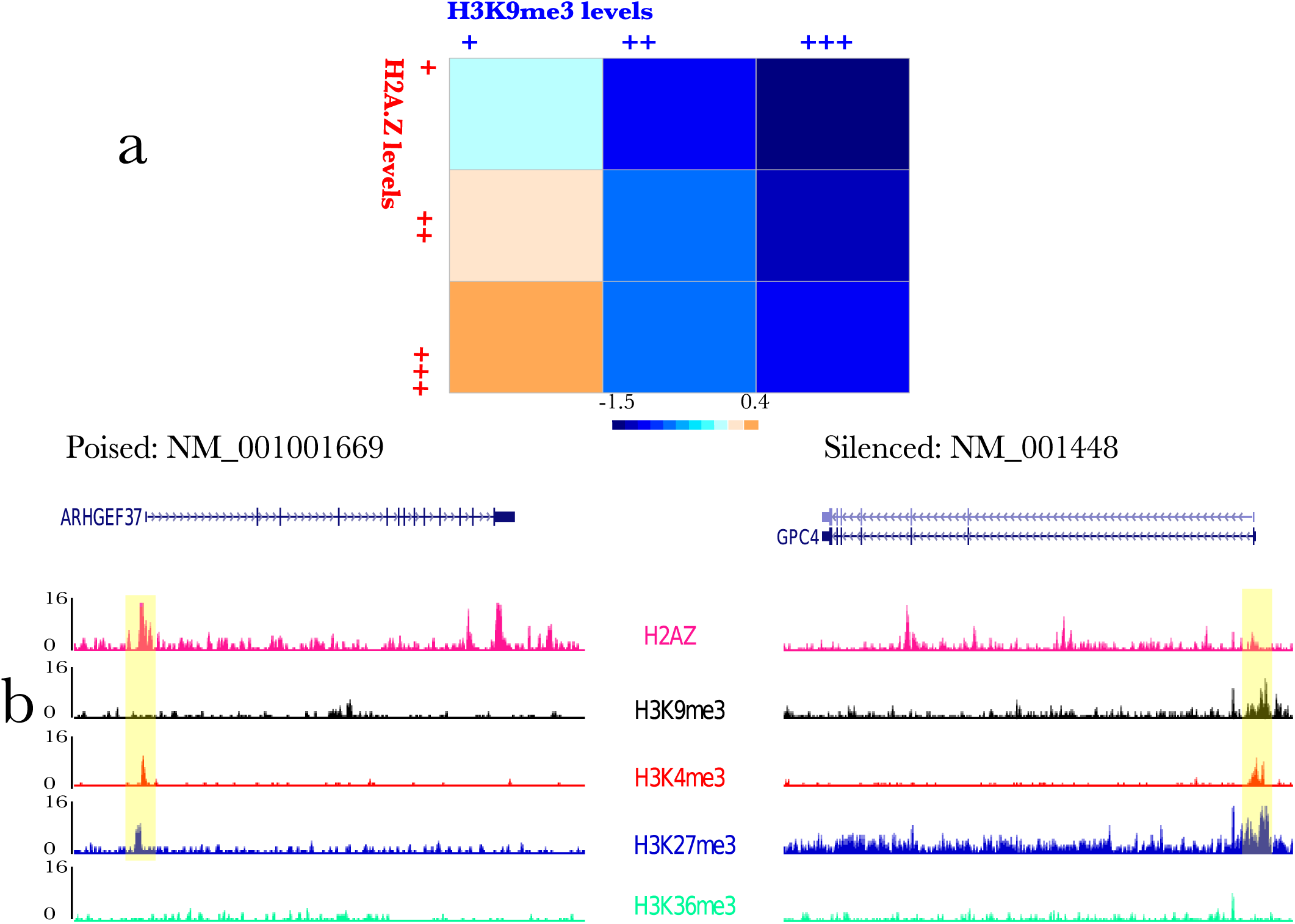
Relative quantities of H2A.Z and H3K9me3 modifications imply genes’ fates during differentiation. The panel (a) is essentially a heat-map, where each square (in total 9 squares) represents a class of genes with a specific combination of H2A.Z and H3K9me3 levels and the color indicates the log-likelihood ratio s (see Materials and Methods for definitions). The panel shows the histone profiles of one poised gene and one silenced gene in HSC as visualized as custom tracks in the UCSC genome browser. Shades in yellow illustrate the promoter regions.

Because a certain genomic region is often co-marked by several types of histone signatures, it would be interesting to examine which levels of other histone marks are associated with poised genes. Based on the HSC dataset, we compared 95 poised genes with the highest levels of H2A.Z and the lowest levels of H3K9me3 (these genes have the largest s values) to 155 silenced genes with the lowest levels of H2A.Z and the highest levels of H3K9me3 (these genes have the smallest s values). As shown in Table 1, these two groups of genes are associated with distinct levels of other histone marks: the poised genes are frequently co-marked by higher levels of H3K27me1, H3K9me1 and H3K4me3, whereas the silenced genes are much less frequently marked by higher levels of these histone marks. In addition, the silenced genes are approximately two times more likely to be co-marked by higher levels of H3K27me3. These observations further confirmed that the relative amount of distinct histone modifications are indicative of poised versus silenced genes.

**Table 1.** High-confidence HSC poised and silenced genes are associated with distinct levels of other histone marks.

|  |  | H3K27<br>me1 | H3K27<br>me3 | H3K36<br>me3 | H3K4<br>me1 | H3K4<br>me3 | H3K9<br>me1 |
| --- | --- | --- | --- | --- | --- | --- | --- |
| Poised<br>(95) | Lev0 | 18 | 59 | 15 | 44 | 9 | 45 |
|  | Lev1+Lev2 | 77 | 36 | 80 | 51 | 86 | 50 |
|  | Proportion <sup>a</sup><br>(%) | 80.1 | 37.9 | 84.2 | 53.7 | 90.5 | 52.6 |
| Silenced<br>(155) | Lev0 | 90 | 45 | 33 | 145 | 114 | 141 |
|  | Lev1+Lev2 | 65 | 110 | 122 | 10 | 41 | 14 |
|  | Proportion<br>(%) | 41.9 | 71.0 | 78.7 | 6.5 | 26.5 | 9.0 |

<sup>a</sup> the proportion of genes with higher levels of modifications (Lev1+ Lev2)

## Discussion

Accumulating data have suggested that histone modifiers and histone marks can pre-set gene expression schemes prior to the differentiation of pluripotent/ multipotent cells [9-11, 19-21]. The prominent “bivalent domain” term has been widely regarded as a mechanism for poising lineage-regulatory genes for expression during the development of ES cells [8]. However, the fate of bivalent genes can be either activation or suppression [8, 11, 12]; in addition, bivalency can also be found in terminally differentiated cells [12, 22]. Therefore, bivalency is not a sufficient or accurate histone signature for poised genes in pluripotent/multipotent cells. In this study, we systematically assessed the poised-gene-discriminating capabilities of various histone marks by taking advantage of quantitative measures. We further proposed a simple probabilistic model for predicting poised genes based only on the H2A.Z and H3K9me3 levels. We showed that our model could be applied to both pluripotent cells and multipotent cells, either human or murine.

H2A.Z is known to be associated with active chromatin, whereas H3K9me3 is involved in gene silencing [11, 23, 24]. Our model illustrates that genes with higher levels of H2A.Z plus lower levels of H3K9me3 are more likely to be poised genes. Thus, it is tempting to argue that an antagonistic outcome between active forces (e.g., H2A.Z) and silencing forces (e.g., H3K9me3) dictates the fate of the corresponding gene during development. Fine-tuning the relative quantities of counter-acting H2A.Z and H3K9me3 histone marks controls the expression potential of a gene.

Our Naïve Bayesian model of H2A.Z and H3K9me3 only shows a modest predictive power, and several factors may have contributed to this result. Firstly, for simplicity, we only considered the histone modification features in promoter regions. However, chromatin has higher-order structure, and the long-range regulation by cis-elements is common[25, 26]. Secondly, as a histone modification pattern may involve multiple constituents, our two-histone mark model might have lacked some information from other histone marks. Nevertheless, we emphasize the wide applicability and simplicity of the model. Lastly, in contrast to the deterministic nature of genetic modulation, epigenetic regulation is much more flexible and plastic; thus, a poised gene becomes activated only when the internal and external environment is permissive.

## Materials and Methods

### Histone modification ChIP-seq raw data collection and preprocessing

The ChIP-seq raw data from CD133^+^ cells (HSCs/ HPCs), CD36^+^ cells (erythrocyte precursors) and GM12878 cells (B lymphoblastoid) were downloaded as FASTQ format files (GEO IDs are GSE12646, GSE36114, GSE29611). The Solexa short reads were mapped to the human genome using Bowtie (index file: hg19.fa) using the default parameters [27]. The human H1 histone modification ChIP-seq data were downloaded as BED files [28], and the mouse ES E14 histone modification ChIP-seq data were downloaded as SRA files (GSE36114, GSE17642). These SRA files were transformed into FASTQ files using the SRA Toolkit from NCBI and then mapped to the mouse genome using Bowtie (index file: mm9.fa) using the default parameters [27]. All of the Bowtie output files were in SAM format, which were further transformed into BED files using a script form MACS [29].

The total reads that fall into a defined genomic region were counted using the intersectBed function from BEDtools [28]. For each cell type, the histone modification tag densities, hereafter referred to as the histone mark feature, were calculated by dividing the read counts by the window sizes. The windows were defined as follows: promoter region (from 2,000 bp upstream to 2,000 bp downstream of TSS, transcription start site) and gene body region (from the TSS upstream 2,000 bp to the TTS, transcription termination site, downstream 2,000 bp). Prior to the Bayesian network analysis and Naïve Bayesian model learning, the histone mark features were split into three nominal variables representing high, moderate, and low levels (termed “Lev2”, “Lev1”, and “Lev0”, respectively). The boundaries between the groups were determined by k-means clustering. 100 repetitions were performed for each histone modification density vector for a given cell type, and the medians were used to define the boundaries.

### Microarray and RNA-seq data sources and analysis

The microarray raw data were downloaded from the GEO database (GEO IDs are GSE12646, GSE26312). Present/absent calls were performed using the MAS5.0 algorithm from the Bioconductor package (http://www.bioconductor.org/). The RNA-seq data of mES E14 were downloaded from the GEO database (GEO ID: GSE36114) in FASTQ format. The raw reads were mapped to the genome using Tophat [30], and transcript abundances were quantitated using Cufflinks [31]. A transcript with an FPKM score over 0.001 was defined as expressed (note that the cut-off may vary according to different sequencing depths and antibodies [32]). As the microarray analysis of the ES cells shows that approximately 1/3 of the total genes are not expressed, this result was used as a guide for setting the cut-off for the present study.

### Naïve Bayesian model and log likelihood ratio s

The core equations of the Naïve Bayesian models in the present study are as follows:

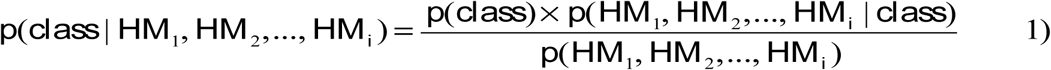

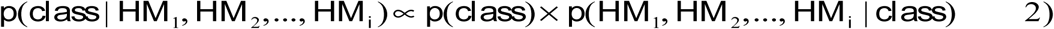

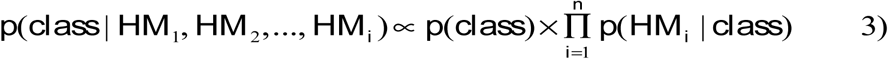

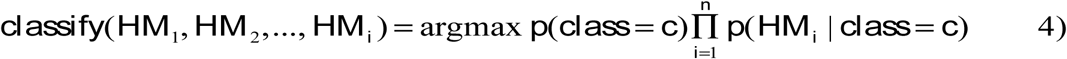

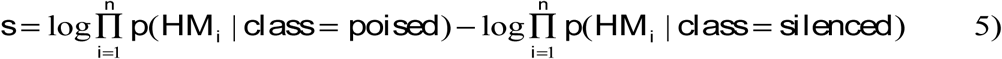

In all these equations, HM_i_ refers to a histone mark profile feature (read density in the promoter region), and class refers to a poised or not poised (silenced) state, as determined by the expression analysis between the progenitor cells and descendant cells. Equation 1) is essentially the Bayesian theorem. In a Naïve Bayesian model under the assumption of independence of all of the predictors (HM_i_), 1) can be deduced to 3). Mathematically, a Naïve Bayesian classifier predicts the instance (i.e., a gene) to be a class that shows the higher joint probability. As there are approximately 20 times more silenced genes compared to poised genes in the HSC dataset, under-sampling of the silenced and poised genes into equal amounts (150 plus 150) was performed. The training of the Naïve Bayesian model was performed using the sampled 150 plus 150 genes, and the remaining genes were used for testing the model. This procedure was repeated 100 times to assess the stability and robustness of the models. The best model was selected from the 100 model based on MCC (Mathew Correlation Coefficient). With regard the modest predictive sensitivity and specificity, when the Naïve Bayesian model learned from the HSC dataset is applied to the other cells, predictions based on original model as specified in equation 4) would be overwhelmed by false positives and false negatives, making the validation of those predictions inapplicable. Therefore, we devised a method to circumvent this problem by defining a log likelihood ratio s, which is intended to rank the probability that a gene is poised versus silenced given its histone mark features. The higher the s value is, the more likely the corresponding gene is poised. As we always train a Naïve Bayesian model by sampling the data into equal amounts of poised and silenced genes, the score s is specified as in equation 5).

## Acknowledgements

The author would like to thank Dr.s Yong Zhang, Xiaobai Zhang, Jianxing Feng, Jinyan Huang, Cheng Li and Qi Liu for very helpful comments and suggestions.

## Additional files

**Additional file 1**

Supp_Figs.ppt is in .ppt format and contains Figures S1-S3. S2 is a scatterplot of poised genes (in red) and silenced genes (in blue) in HSC. Varying sizes of dots correspond to H3K36me3 levels, the bigger the size is, the higher the H3K36me3 level is.

## References

1. Heintzman ND, Hon GC, Hawkins RD, Kheradpour P, Stark A, Harp LF, Ye Z, Lee LK, Stuart RK, Ching CW et al: Histone modifications at human enhancers reflect global cell-type-specific gene expression. Nature 2009, 459(7243):108–112.

2. Zhao Q, Zhang Y: Epigenome sequencing comes of age in development, differentiation and disease mechanism research. Epigenomics 2011, 3(2):207–220.

3. Barski A, Jothi R, Cuddapah S, Cui K, Roh TY, Schones DE, Zhao K: Chromatin poises miRNA- and protein-coding genes for expression. Genome Res 2009, 19(10):1742–1751.

4. Wang Z, Zang C, Rosenfeld JA, Schones DE, Barski A, Cuddapah S, Cui K, Roh TY, Peng W, Zhang MQ et al: Combinatorial patterns of histone acetylations and methylations in the human genome. Nat Genet 2008, 40(7):897–903.

5. Tian R, Feng J, Cai X, Zhang Y: Local chromatin dynamics of transcription factors imply cell-lineage specific functions during cellular differentiation. Epigenetics 2012, 7(1).

6. He HH, Meyer CA, Shin H, Bailey ST, Wei G, Wang Q, Zhang Y, Xu K, Ni M, Lupien M et al: Nucleosome dynamics define transcriptional enhancers. Nat Genet 2010, 42(4):343–347.

7. Zhang Z, Zhang MQ: Histone modification profiles are predictive for tissue/cell-type specific expression of both protein-coding and microRNA genes. BMC Bioinformatics 2011, 12:155.

8. Bernstein BE, Mikkelsen TS, Xie X, Kamal M, Huebert DJ, Cuff J, Fry B, Meissner A, Wernig M, Plath K et al: A bivalent chromatin structure marks key developmental genes in embryonic stem cells. Cell 2006, 125(2):315– 326.

9. Lindeman LC, Andersen IS, Reiner AH, Li N, Aanes H, Ostrup O, Winata C, Mathavan S, Muller F, Alestrom P et al: Prepatterning of developmental gene expression by modified histones before zygotic genome activation. Dev Cell 2011, 21(6):993–1004.

10. Xu CR, Cole PA, Meyers DJ, Kormish J, Dent S, Zaret KS: Chromatin “prepattern” and histone modifiers in a fate choice for liver and pancreas. Science 2011, 332(6032):963–966.

11. Cui K, Zang C, Roh TY, Schones DE, Childs RW, Peng W, Zhao K: Chromatin signatures in multipotent human hematopoietic stem cells indicate the fate of bivalent genes during differentiation. Cell Stem Cell 2009, 4(1):80–93.

12. De Gobbi M, Garrick D, Lynch M, Vernimmen D, Hughes JR, Goardon N, Luc S, Lower KM, Sloane-Stanley JA, Pina C et al: Generation of bivalent chromatin domains during cell fate decisions. Epigenetics Chromatin 2011, 4(1):9.

13. Ansel KM, Djuretic I, Tanasa B, Rao A: Regulation of Th2 differentiation and Il4 locus accessibility. Annu Rev Immunol 2006, 24:607-656.

14. de Laat W, Klous P, Kooren J, Noordermeer D, Palstra RJ, Simonis M, Splinter E, Grosveld F: Three-dimensional organization of gene expression in erythroid cells. Curr Top Dev Biol 2008, 82:117–139.

15. Combining SVMs with Various Feature Selection Strategies [http://www.csie.ntu.edu.tw/∼cjlin/papers/features.pdf]

16. e1071: Misc Functions of the Department of Statistics (e1071) [ http://cran.R-project.org]

17. Yu H, Zhu S, Zhou B, Xue H, Han JD: Inferring causal relationships among different histone modifications and gene expression. Genome Res 2008, 18(8):1314–1324.

18. Chickering DM: The WinMine toolkit. In: Microsoft Research Technical Report MSR-TR-2002-103. Redmond, WA: Microsoft; 2002.

19. Guenther MG, Levine SS, Boyer LA, Jaenisch R, Young RA: A chromatin landmark and transcription initiation at most promoters in human cells. Cell 2007, 130(1):77–88.

20. Blythe SA, Klein PS: Prepatterning embryonic development: tabula scripta? Dev Cell 2011, 21(6):977–978.

21. Hong SH, Rampalli S, Lee JB, McNicol J, Collins T, Draper JS, Bhatia M: Cell fate potential of human pluripotent stem cells is encoded by histone modifications. Cell Stem Cell 2011, 9(1):24–36.

22. Roh TY, Cuddapah S, Cui K, Zhao K: The genomic landscape of histone modifications in human T cells. Proc Natl Acad Sci U S A 2006, 103(43):15782–15787.

23. Creyghton MP, Markoulaki S, Levine SS, Hanna J, Lodato MA, Sha K, Young RA, Jaenisch R, Boyer LA: H2AZ is enriched at polycomb complex target genes in ES cells and is necessary for lineage commitment. Cell 2008, 135(4):649–661.

24. Rosenfeld JA, Xuan Z, DeSalle R: Investigating repetitively matching short sequencing reads: the enigmatic nature of H3K9me3. Epigenetics 2009, 4(7):476–486.

25. Lieberman-Aiden E, van Berkum NL, Williams L, Imakaev M, Ragoczy T, Telling A, Amit I, Lajoie BR, Sabo PJ, Dorschner MO et al: Comprehensive mapping of long-range interactions reveals folding principles of the human genome. Science 2009, 326(5950):289–293.

26. Liu L, Zhang, Y., Feng, J., Zheng, N.,Yin, J., Zhang, Y.: GeSICA: Genome Segmentation from Intra Chromosomal Associations. In: BMC Genomics (in press). 2012.

27. Langmead B, Trapnell C, Pop M, Salzberg SL: Ultrafast and memory-efficient alignment of short DNA sequences to the human genome. Genome Biol 2009, 10(3):R25.

28. Quinlan AR, Hall IM: BEDTools: a flexible suite of utilities for comparing genomic features. Bioinformatics 2010, 26(6):841–842.

29. Zhang Y, Liu T, Meyer CA, Eeckhoute J, Johnson DS, Bernstein BE, Nusbaum C, Myers RM, Brown M, Li W et al: Model-based analysis of ChIP-Seq (MACS). Genome Biol 2008, 9(9):R137.

30. Trapnell C, Pachter L, Salzberg SL: TopHat: discovering splice junctions with RNA-Seq. Bioinformatics 2009, 25(9):1105–1111.

31. Trapnell C, Roberts A, Goff L, Pertea G, Kim D, Kelley DR, Pimentel H, Salzberg SL, Rinn JL, Pachter L: Differential gene and transcript expression analysis of RNA-seq experiments with TopHat and Cufflinks. Nat Protoc 2012, 7(3):562–578.

32. Toung JM, Morley M, Li MY, Cheung VG: RNA-sequence analysis of human B-cells. Genome Research 2011, 21(6):991–998.

